# Positively and negatively autocorrelated environmental fluctuations have opposing effects on species coexistence

**DOI:** 10.1101/869073

**Authors:** Sebastian J. Schreiber

## Abstract

Environmental fluctuations can mediate coexistence between competing species via the storage effect. This fluctuation-dependent coexistence mechanism requires three conditions: (i) a positive covariance between environment conditions and the strength of competition, (ii) species-specific environmental responses, and (iii) species are less sensitive to competition in environmentally unfavorable years. In serially uncorrelated environments, condition (i) only occurs if favorable environmental conditions immediately and directly increase the strength of competition. For many demographic parameters, this direct link between favorable years and competition may not exist. Moreover, many environmental variables are temporal autocorrelated, but theory has largely focused on serially uncorrelated environments. To address this gap, a model of competing species in autocorrelated environments is analyzed. This analysis shows that positive autocorrelations in demographic rates that increase fitness (e.g. maximal fecundity or adult survival) produce the positive environment-competition covariance. Hence, when these demographic rates contribute to buffered population growth, positive temporal autocorrelations generate a storage effect, otherwise they destabilize competitive interactions. For negatively autocorrelated environments, this theory highlights an alternative stabilizing mechanism that requires three conditions: (i’) a negative environmental-competition covariance, (ii) species-specific environmental responses, and (iii’) species are less sensitive to competition in more favorable years. When the conditions for either of these stabilizing mechanisms are violated, temporal autocorrelations can generate stochastic priority effects or hasten competitive exclusion. Collectively, these results highlight that temporal autocorrelations in environmental conditions can play a fundamental role in determining ecological outcomes of competing species.

## Introduction

Most species are engaged in competitive interactions with other species [Gurevitch et al., 1992, Kaplan and Denno, 2007]. This mutual antagonism can result in one species driving other species extinct. According to ecological theory, this competitive exclusion is inevitable when there are more species than limiting factors and the community approaches a steady state [Volterra, 1928, McGehee and Armstrong, 1977]. Hutchinson [1961] proposed that fluctuations in environmental conditions may allow species competing for few limiting factors to coexist. In an series of influential papers [Chesson and Warner, 1981, Chesson, 1983, 1988, 1994], Peter Chesson developed a mathematical theory for when and how environmental fluctuations, via nonlinear averaging and the storage effect, mediate species coexistence. In the past decade, empirical evidence for these coexistence mechanisms were identified in a diversity of systems including zooplankton [Cáceres, 1997], prairie grasses [Adler et al., 2006], desert annual plants [Angert et al., 2009], tropical trees [Usinowicz et al., 2012], phytoplankton [Ellner et al., 2016], sagebrush [Chu and Adler, 2015, Ellner et al., 2016], and nectar yeasts [Letten et al., 2018].

Until recently [Benaïm and Schreiber, 2019], mathematical methods for studying coexistence for species experiencing environmental stochasticity assumed these fluctuations are uncorrelated in time [Chesson, 1982, Chesson and Ellner, 1989, Ellner, 1989, Schreiber et al., 2011, Hening and Nguyen, 2018], and ecological theory has mostly focused on this case [Chesson, 1994, 2000, Angert et al., 2009, Stump and Chesson, 2017, Kortessis and Chesson, 2019]. Environmental fluctuations, however, are often autocorrelated [Steele, 1985]. Minimum and maximal monthly temperatures in both terrestrial and marine systems are typically positively autocorrelated [Vasseur and Yodzis, 2004]; months with higher temperature maxima tend to be followed by months with higher maxima. Approximately 20% of terrestrial sites on earth exhibit positively autocorrelated yearly rainfall, while 5% exhibit negatively autocorrelated rainfall [Sun et al., 2018]. Although considered less frequently, negative autocorrelations may be common in other situations [Metcalf and Koons, 2007]. For example, density-dependent driven oscillations of an herbivore or predator can result in negatively autocorrelated fluctuations in the mortality rates of its prey species.

Theoretical and empirical studies show that autocorrelated, environmental fluctuations can have large impacts on population demography [Foley, 1994, Petchey et al., 1997, Cuddington and Yodzis, 1999, Pike et al., 2004, Gonzalez and Holt, 2002, Cuddington and Hastings, 2016]. Theory predicts that positively autocorrelated fluctuations increase extinction risk when populations exhibit under-compensatory dynamics, but decreases extinction risk when populations exhibit overcompensatory dynamics [Petchey et al., 1997]. Consistent with the first theoretical prediction, clonal populations of *Folsomia candida* exhibited shorter times to extinction when fluctuating mortality rates were positively autocorrelated [Pike et al., 2004]. For structured populations, temporal autocorrelations can alter longterm population growth rates [Roy et al., 2005, Tuljapurkar and Haridas, 2006, Schreiber, 2010]. For example, lab experiments with paramecia and theory predicted that positively autocorrelated fluctuations in the local fitnesses of spatially structured populations can increase long-term population growth rates [Roy et al., 2005, Matthews and Gonzalez, 2007, Schreiber, 2010].

As autocorrelated fluctuations are common and have demographic impacts, they likely influence ecological outcomes of competing species. As recent mathematical theory [Benaïm and Schreiber, 2019] provides the analytical tools to explore this influence, I analyze models of two species competition accounting for autocorrelated fluctuations in environmental conditions. As these models correspond to competition for a single, limiting factor and undercompensatory density-dependence, stable coexistence doesn’t occur in constant environments, only neutral coexistence is possible. Moreover, uncorrelated environmental fluctuations do not stabilize neutral coexistence in these models. Therefore, using a mixture of analytical and numerical approaches applied to the stochastic models, I tackle the following questions: When do positively or negatively autocorrelated fluctuations mediate coexistence? When do they disrupt neutral coexistence and, if they do, is the identity of the excluded species predictable? What types of shifts in competitive outcomes are possible as temporal autocorrelations shift from negative to positive?

## Model and Methods

Consider two competing species with densities *n*_1_ and *n*_2_. The fitness of individuals within species *i, f* (*C, E*_*i*_), decreases with the strength of competition *C* and increases with respect to an environmental response variable *E*_*i*_. The strength of competition is given by a weighted combination of the species densities, *C* = *a*_1_*n*_1_ + *a*_2_*n*_2_ where *a*_*i*_ determines the per-capita contribution of species *i* to the strength of competition. The environmental variable *E*_*i*_ represents the net effect of environmental conditions on species *i*’s fitness. Unlike prior work on the storage effect [e.g. Chesson, 1994, Kuang and Chesson, 2009, Stump and Chesson, 2017], the strength of competition *C* is not a function of the environmental variables *E*_*i*_. Therefore, for uncorrelated environmental fluctuations, there is no covariance between the environment and the strength of competition and, consequently, no storage effect [Chesson, 1994]. Consistent with meteorological models of various weather variables [Wilks and Wilby, 1999, Semenov, 2008], fluctuations in the environmental response variables follow a multivariate autoregressive (MAR) process with means 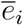, standard deviations *σ*_*i*_, cross-correlation *τ*, and temporal autocorrelation *ρ*. Under these assumptions, the dynamics of the system are

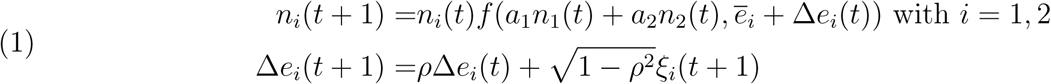

where Δ*e*_*i*_(*t*) are the deviations of the environmental variables away from the means 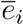, *ξ*_*i*_(*t*) are independent in time with mean 0, standard deviation *σ*_*i*_, and cross-correlation Corr[*ξ*_1_(*t*)*ξ*_2_(*t*)] = *τ*. Provided that |*ρ*| < 1 (i.e. the environmental fluctuations are neither perfectly positively or negatively autocorrelated), the environmental deviations Δ*e*_*i*_(*t*) converge to a unique stationary distribution 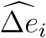 with mean 0, variance 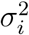, and cross-correlation *τ*. The 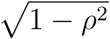 term in (1) allows one to independently vary the variance of the stationary distribution and the temporal autocorrelation.

I study the dynamics of (1) using a mixture of analytical and numerical methods. The analysis is based the per-capita growth rates log *f* averaged over fluctuations in the environmental variables and the strength of competition [Chesson, 1994]. The analysis assumes that the maximal per-capita growth rates, 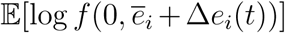, of each species in the absence of competition are positive, and the fitness function *f* exhibits compensating density-dependence. These assumptions ensure there exists a unique, positive stationary distributions 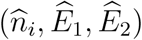 for species *i* in the absence of competition with species *j* ≠ *i* [Benaïm and Schreiber, 2009]. Here, 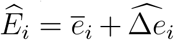 and the stationary distribution of the competitive strengths equals 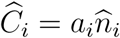. Conditions for coexistence or exclusion depend on the invasion growth rate of species *j* ≠ *i*

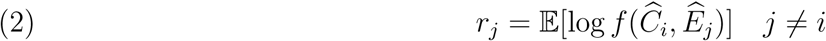

that corresponds to the average per-capita growth rate of species *j* when it is rare and species *i* is common. If both invasion growth rates are positive (*r*_1_ > 0, *r*_2_ > 0), then the species coexist in the sense of stochastic persistence [cf. Theorem 1 in Benaïm and Schreiber, 2019] i.e. a statistical tendency to stay away from the extinction set (Fig. 1A). If *r*_1_ < 0 < *r*_2_ (respectively, *r*_2_ < 0 < *r*_1_), then species 2 excludes species 1 (respectively, species 1 excludes species 2) [cf. of Corollary 2 in Benaïm and Schreiber, 2019] (Fig. 1B). Finally, if both invasion growth rates are negative, then the system exhibits a stochastic bistability: whenever both species are initially present, there is a positive probability (*p* > 0 depending on the initial densities) that species 1 is excluded, and a complementary, positive probability (1 − *p* > 0) that species 2 is excluded [cf. of Theorem 3 and Corollary 2 in Benaïm and Schreiber, 2019].

**Figure 1.**
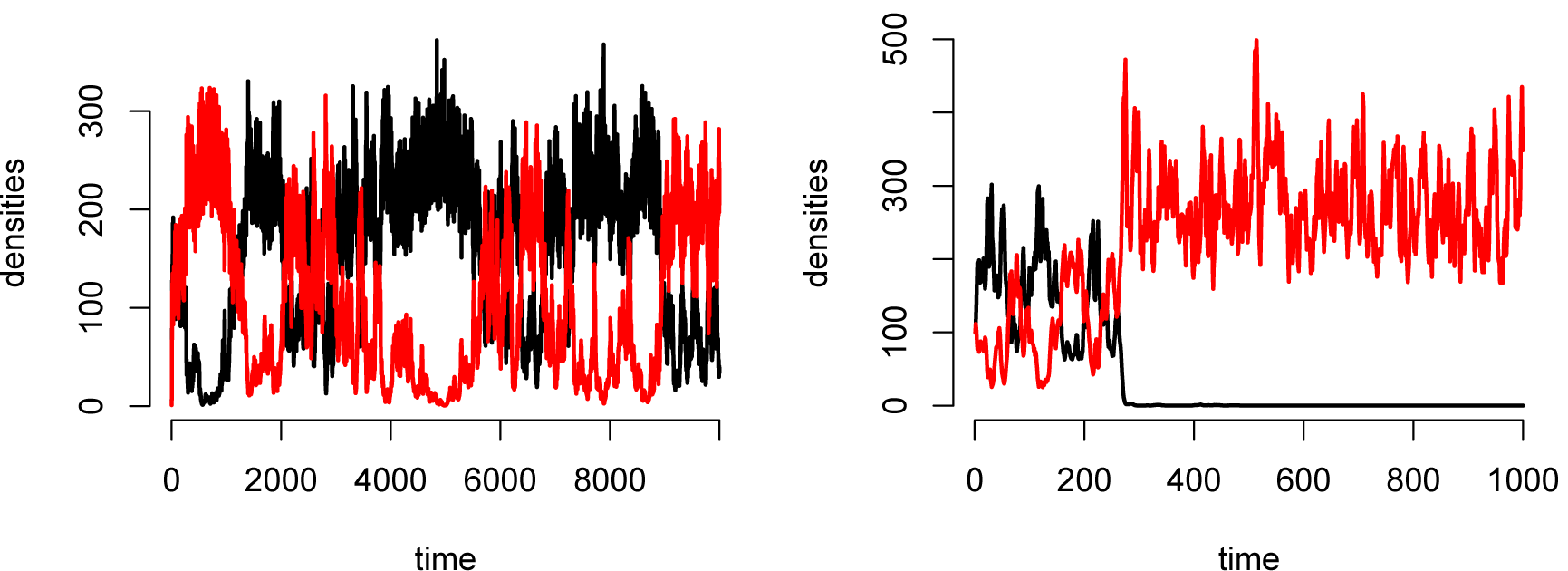
Stochastic coexistence (A) or exclusion (B) for competition in a fluctuating environment. Parameters: The fitness function (4) with fluctuations in survival, with λ = 2, *a*_1_ = *a*_2_ = 0.01. *ξ*_*i*_(*t*) are normally distributed with means 0, standard deviations *σ*_*i*_ = 0.5, cross-correlation *τ* = 1, and autocorrelations *ρ* = 0.5 in (A) and *ρ* =0.5 in (B).

To derive analytic approximations for the invasion growth rates *r*_*j*_, I use a diffusion-type scaling in which 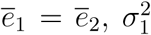, and 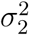 are small and of the same order [Turelli, 1977, Karlin and Taylor, 1981]. To illustrate the analytic results for the general model (1), I numerically compute the invasion growth rates for fitness functions *f* with a density-dependent component λ*/*(1 + *C*) and a density-independent component *s*. In the theoretical and empirical literature [e.g., Adler et al., 2007, Godoy et al., 2014], the density-dependent term, usually, is interpreted as a maximal per-capita fecundity λ that is reduced by competition *C*, while the density-independent term 0 < *s* < 1 corresponds to survivorship. I examine environmentally driven fluctuations in both terms. For fluctuations in the density-dependent term, the fitness function equals

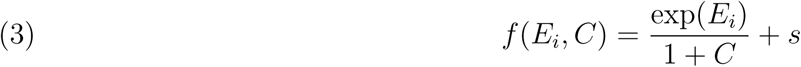

while for fluctuations in the density-independent terms, it equals

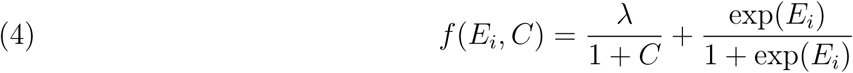

In both cases, the variables *ξ*_*i*_(*t*) are drawn from bivariate normals with mean 0, variances 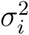, and cross-correlation *τ*. Thus, with their traditional interpretation, the fluctuations in (3) correspond to lognormally distributed fluctuations in maximal fecundities, and the fluctuations in (4) correspond to logit-normally distributed fluctuations in survival. While I focus on this traditional interpretation of (3) and (4), one can reverse their interpretation for species where density-dependence acts more strongly on survivorship than reproduction, for example in salmonids [Grossman and Simon, 2020].

## Results

I first present results for the deterministic model that show stable coexistence does not occur without environmental fluctuations. Next, I present results for environmental fluctuations where species only differ to the degree their environmental responses are correlated. In this special case, the deterministic dynamics are neutrally stable and environmental fluctuations either lead to coexistence, fluctuating neutral coexistence, or to a stochastic priority effect. Finally, I present results in which the species differ in their mean environmental response 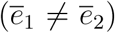, the variation in their environmental responses (*σ*_1_ ≠ *σ*_2_), and the correlation in their environmental responses. These results highlight when environmental stochasticity reverses competitive outcomes as well as stabilizes competitive interactions.

### Neutrality or exclusion in constant environments

When the mean environmental responses are equal 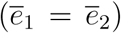 and there are no environmental fluctuations (*σ*_1_ = *σ*_2_ = 0), the species exhibit neutral coexistence (proof in Appendix). Specifically, there exists a line of equilibria connecting the single species equilibria. Community trajectories always converge to one of these equilibria, but different initial conditions can converge to different equilibria. This coexistence isn’t stable in that small pulse perturbations typically shift the community to a different equilibrium state [Schreiber, 2006]. In contrast, if one species has a higher mean environmental response than the other 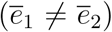, then this species competitively excludes the other species (proof in Appendix). These conclusions are consistent with general theory on limiting similarity [see, e.g., Meszéna et al., 2006, Pasztor et al., 2016]

### From Neutrality to Coexistence or Alternative Stable States

The simplest case in which environmental fluctuations alter ecological outcomes is when both species have the same mean environmental response 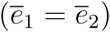 and experience the same degree of variation in their environmental response (*σ*_1_ = *σ*_2_). Without environmental fluctuations, this leads to neutral coexistence in which *r*_*j*_ = 0 for both species. With environmental fluctuations, our diffusion approximation for model (1) yields (derivation in Appendix)

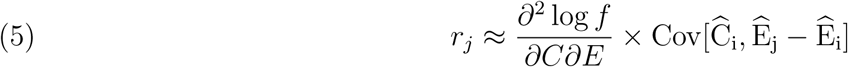

where the mixed partial derivative, 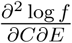, is evaluated at 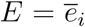 and the equilibrium value 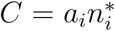 where 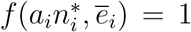. The sign of this mixed partial derivative determines whether the log fitness function is superadditive (positive sign) or subadditive (negative sign) with respect to the interactive effects of competition and environmental fluctuations [see e.g. Puterman, 2014]. Subadditivity means that the log fitness function, log *f*, is less sensitive to the effects of competition when environmental conditions are poor. This corresponds to population buffering, a necessary component of the storage effect [Chesson, 1994, Ellner et al., 2016]. Superadditivity, in contrast, means that the log fitness function is more sensitive to the effects of competition when environmental conditions are poor, but also that is less sensitive to the effects of competition when environmental conditions are good. For example, for the fitness function (3) with fluctuations in fecundity and positive survival 0 < *s* < 1, the log-fitness function is subadditive and population buffering occurs. In contrast, for the fitness function (4) with fluctuations in survival, the log fitness function is superadditive and populations are less sensitive to competition in years with high survival.

The final term in equation (5) corresponds to the covariance between the strength of competition due to the common species (*C*_*i*_) and the difference (*E*_*j*_ − *E*_*i*_) between the environmental responses of the rare and common species. This term is positive when years with high and low densities, respectively, of the common species also correspond to years where the rare species has a higher and lower environmental response, respectively. This covariance for model (1) is proportional to a product of three terms (derivation in Appendix):

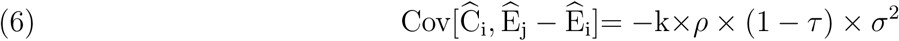

where the proportionality term *k* = *k*(*ρ*) is positive for all *ρ* and *k*(*ρ*) × *ρ* is an increasing function of *ρ*. When the environmental fluctuations are serially uncorrelated (*ρ* = 0) or the species have identical responses to the environment (*τ* = 1), this covariance is zero and, consequently, the invasion growth rates equal 0, and the species exhibit a fluctuating form of neutral coexistence. In contrast, when there are species-specific response to the environment (*τ* < 1) and environmental fluctuations are autocorrelated (*ρ* ≠ 0), equation (6) implies that the sign of 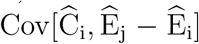 is opposite of the sign of the autocorrelation. Hence, in positively autocorrelated environments, years with greater densities of the common species tend to be years where the environmental conditions are less favorable to the rare species. Moreover, the magnitude of the covariance is greater for positive autocorrelations than negative autocorrelations i.e. 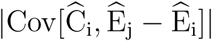 is greater for a given *ρ* > 0 than for the corresponding − *ρ* < 0 (see Appendix for derivation). Intuitively, positive autocorrelations generate greater variance in the strength of competition than negative autocorrelations and, thereby, yield a greater covariance. Collectively, equations (5) and (6) imply environmental fluctuations promote coexistence (i.e. *r*_*j*_ > 0) in two situations: (i) the log-fitness is superadditive and environmental fluctuations are negatively autocorrelated, and (ii) the log-fitness is subadditive and environmental fluctuations are positively autocorrelated i.e. the storage effect. In contrast, if (iii) the log-fitness function is superadditive and environmental functions are positive autocorrelated or (iv) the log-fitness function is subadditive and the fluctuations are negatively autocorrelated, then the system is stochastically bistable: with complementary, positive probabilities species 1 or 2 is excluded.

Figure 2 illustrates these analytical conclusions for the fitness function (3) with fluctuating fecundity or (4) with fluctuating survival. As the log fitness for (3) is subadditive with respect to fecundity, positively autocorrelated fluctuations in fecundity mediate coexistence, while negative autocorrelations lead to stochastic bistability (Fig. 2A). In contrast, as the log fitness for (4) is superadditive with respect to survival, positively autocorrelated fluctuations in survival lead to stochastic bistability, while negative autocorrelations promote coexistence (Fig. 2B).

**Figure 2.**
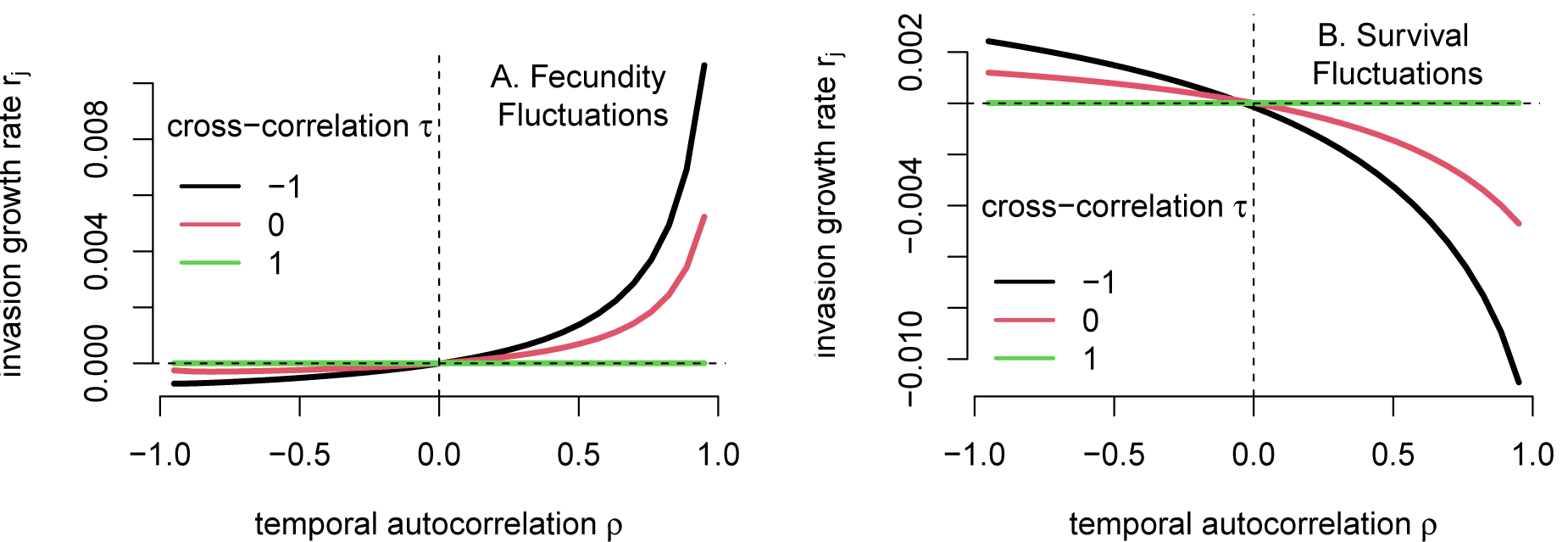
Positively autocorrelated fluctuations in fecundity (A) and negative autocorrelated fluctuations in survival (B) mediate coexistence. Numerically computed invasion growth rates *r*_*j*_ as a function of the temporal autocorrelation *ρ* for competing species. In (A), the fitness function *f* is given by (3) and there are fluctuations in fecundity. In (B), the fitness function is given by (4) with fluctuations in survival. Different curves corresponds to different levels of cross-correlation *τ* where competitors only differ demographically if *τ* < 1. For positive invasion growth rates, the competitors coexist (stochastic persistence). For negative invasion growth rates, each competitor is excluded with positive complementary positive probabilities (stochastic bistability). Parameters: *a*_1_ = *a*_2_ = 1 and (*ξ*_1_(*t*), *ξ*_2_(*t*)) is normally distributed with standard deviation 0.3 for both panels. For A, *s* = 0.9, and 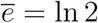. For B, 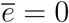, and λ = 2.

### The Effects of Fitness Differences and Nonlinear Averaging

Asymmetries in the mean response 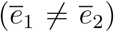 and the variability of these responses (*σ*_1_ ≠ *σ*_2_) lead to two additional terms in the invasion growth rate for model (1) (derivation in Appendix):

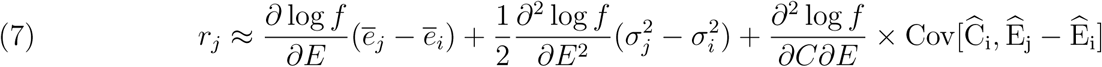

**Figure 3.**
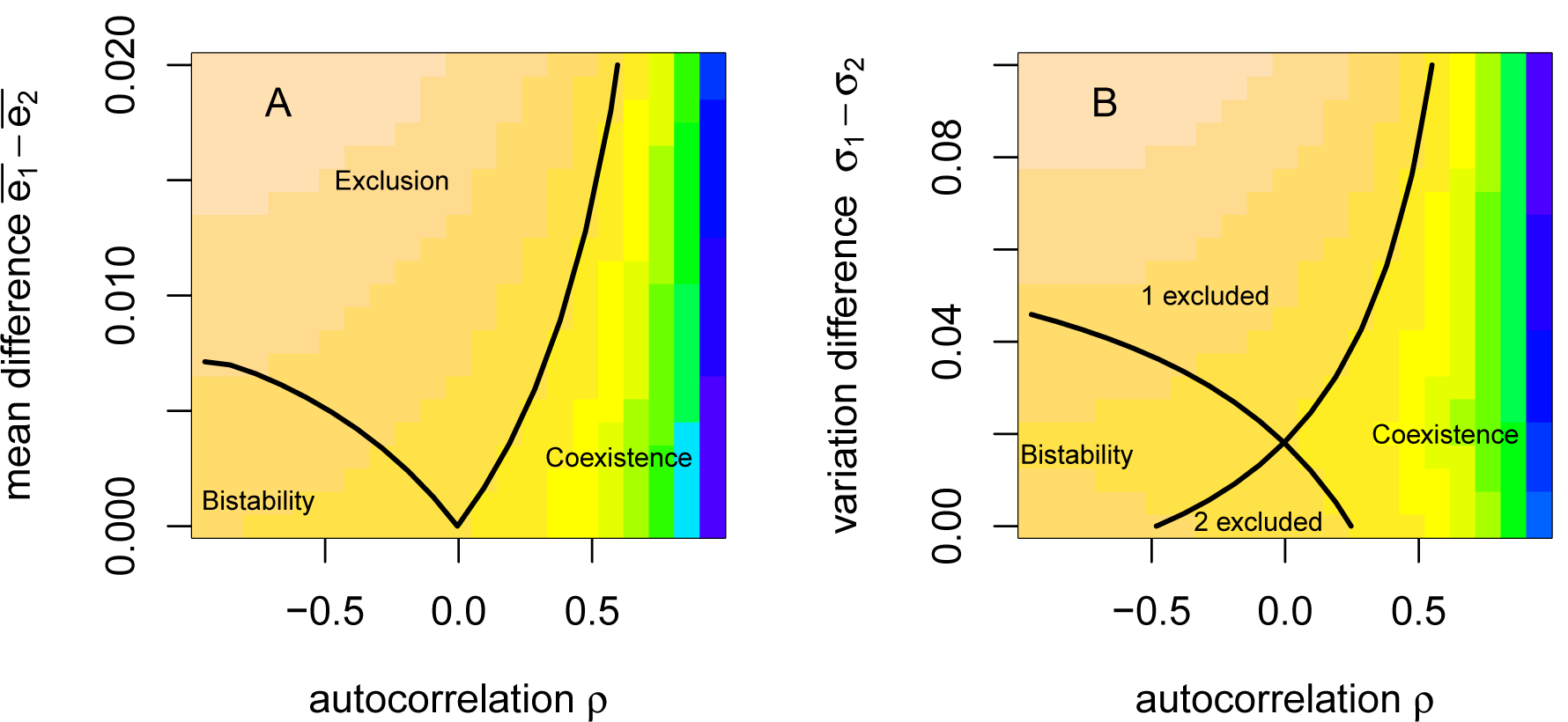
Interactive effects of fitness differences and autocorrelation on ecological outcomes. In A, species 1 has a larger mean environmental response 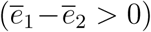 over species 2. In B, species 1 has a slightly stronger mean environmental response than species 2 but experiences greater environmental variation (*σ*_1_ − *σ*_2_ ≥ 0) than species 2. Regions of coexistence, competitive exclusion, and bistability are shown. Solid contour lines correspond to *r*_1_ = 0 and *r*_2_ = 0. Coloring is determined by the minimum of the invasion growth rates, min *r*_1_, *r*_2_, with tan for most negative values to blue for the most positive values. Parameters: fitness function *f* is given by (3) with 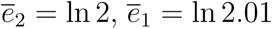 in B, 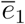 as determined by the mean difference in the environmental response 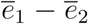 in A, *ξ*_*i*_(*t*) are normally distributed with *τ* = −1, *σ*_1_ = *σ*_2_ = 0.3 in A, and *σ*_2_ = 0.3 and *σ*_1_ as shown in B.

As the log fitness increases with the environmental response variable (*E*), the first term in (7) is proportional to the difference, 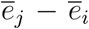, in the mean environmental response between the rare and common species. Intuitively, when the rare species benefits more, on average, from the environmental conditions 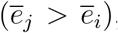, its invasion growth rate is larger. The sign of the second term in equation (7) depends on the concavity of log fitness with respect to the environmental response variable at 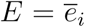 and 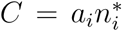. When the log fitness function is concave at this point, the second term contributes positively to the invasion growth if the rare species exhibits less variation in its environmental response 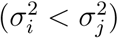. This second term corresponds to the effect of nonlinear averaging [Chesson, 1994]. Namely, there is a reduction (respectively, increase) in the invasion growth rate due to the concavity (respectively, convexity) of the log fitness function at 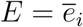 and 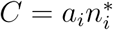.

Differences in the variation of the environmental responses also impact the covariance between the density of common species and difference in the environmental responses i.e. the third term of (7). Specifically, a refinement of expression (6) shows that (derivation in Appendix)

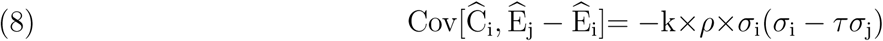

To better understand this expression, I partition it into the sum of two terms: a community average term and a species-specific term [Chesson, 2018]. The community average term equals the average of the covariances 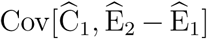 and 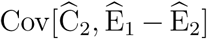 and contributes equally to both species invasion growth rates. This community average equals

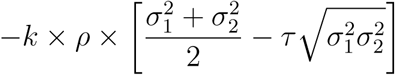

As the the arithmetic mean 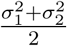 is greater than the geometric mean 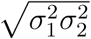, the community average term has the opposite sign of the sign of the autocorrelation *ρ*. On the other hand, the species-specific term equals

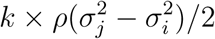

Thus, the species-specific contribution to 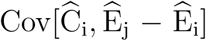 has the same sign as the autocorrelation *ρ* for the species with larger variance, and the opposite sign for the species with smaller variance. This asymmetry may benefit or harm the species with the smaller variance. For example, if there is population buffering and positively autocorrelated fluctuations, then the species with the smaller variance benefits more from the storage effect than the species with the larger variance. Alternatively, if there is negative population buffering and negative autocorrelated fluctuations, then the species with the larger variance benefits more from the environment-competition covariance.

To illustrate these analytical results, Fig. 3 numerically computed the invasion growth rates *r*_*j*_ with the fitness function (3) with fluctuations in fecundity. In Fig. 3A, the species only exhibit differences in their mean environmental response and have negatively correlated environmental responses. With small differences in the mean environmental response, sufficiently negative autocorrelations result in bistability, intermediate autocorrelations result in the species with larger mean environmental response excluding the other species, and sufficiently positive autocorrelations mediate coexistence. When the difference in mean environmental response is large, sufficiently positive autocorrelations mediate coexistence, otherwise the species with the lower mean environmental response is excluded. In Fig. 3B, one species has a higher mean environmental response 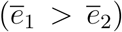, but also experiences greater variability in its environmental response (*σ*_1_ > *σ*_2_). When the difference in variation is sufficiently low, species 1 excludes species 2 unless temporal autocorrelation is sufficiently high. In contrast, when the difference in variation is sufficiently high, species 2 excludes species 1 unless temporal autocorrelation is sufficiently high. This reversal in fates stems from the reduction of *r*_1_ due to greater environmental variation simultaneously increasing the negative effect of nonlinear averaging and reducing the strength of the storage effect.

## Discussion

Hutchinson [1961] wrote “the diversity of the plankton was explicable primarily by a permanent failure to achieve equilibrium as the relevant external factors changes.” It wasn’t until thirty years later that Peter Chesson developed a theoretical framework for precisely identifying fluctuation-dependent mechanisms for coexistence [see, e.g., Chesson and Warner, 1981, Chesson, 1983, 1988, 1994] including the storage effect. The storage effect stabilizes coexistence, when (i) there is a positive correlation between the environmental response of each species and the competition experienced by that species, (ii) there are species-specific environmental responses, and (iii) buffered population growth in which species are less sensitive to competition in years of poor environmental conditions. In serially uncorrelated environments, the first condition requires that there is a direct and immediate impact of the environmental response on the strength of competition. This occurs, for example, in annual populations with year to year variation in germination rates: when more seeds germinate, more plants compete for limiting resources. This direct and immediate impact, however, does not occur when maximal yield or adult survival varies, as in the models considered here. However, when the temporal fluctuations in these demographic rates are autocorrelated, the analysis presented here reveals that the environmental-competition covariance can still occur in these fundamental demographic parameters. Furthermore, these autocorrelated fluctuations highlight how another, under-appreciated stabilizing mechanisms arises when conditions (i) and (iii) simultaneously do not hold.

Hutchinson [1961] hinted at temporal autocorrelations as a stabilizing mechanism when he wrote: “equilibrium would never be expected in nature whenever complete competitive replacement of one species by another occurred in a time (*t*_*c*_), of the same order, as the time (*t*_*e*_) taken for a significant seasonal change in the environment.” As competitive exclusion typically takes several generations, Hutchinson’s quote implies that coexistence requires the shifts in environmental conditions favoring differing species should take several generations. Thus, the environmental conditions must be positively autocorrelated over several generations. Consistent with this suggestion, I show that when there is population buffering (i.e. log fitness is subadditive) and there are species-specific environmental responses, positively autocorrelated fluctuations in environmental conditions yield a storage effect. Namely, condition (i) is satisfied as better years for one species tend to be preceded by better years for this species and, therefore, tends to lead to higher densities (greater competition) in the focal year. The strength of this environment-competition covariance depends on the degree that competitive communities are biotically saturated with individuals. For saturated communities, like those modeled by Chesson’s lottery model [Chesson and Warner, 1981] and Hubbell’s neutral model [Hubbell, 2001], autocorrelated fluctuations can not generate this covariance when one competitor is rare; the abundance of the common species remains relatively constant. In contrast, for less saturated communities, which are common [Houlahan et al., 2007], fluctuations in environmental conditions can lead to fluctuations in the abundance of the common species and, when positively autocorrelated, lead to a positive environment-competition covariance. Our findings about positive autocorrelations generating a storage effect are consistent with two prior studies [Jiang and Morin, 2007, Schreiber et al., 2019]. Jiang and Morin [2007] manipulated temporal fluctuations experienced by two species of ciliated protists competing for bacterial resources. These temperature fluctuations had large effects on the intrinsic growth rates of the two species, consistent with the fluctuating fecundity model considered here. When the temperature fluctuations were temporally uncorrelated, their experimental results suggested that resource partitioning and temperature-dependent competitive effects lead to coexistence, not a storage effect. In contrast, when temperature fluctuations were positively autocorrelated, their experimental results suggested that resource partitioning and the storage effect lead to coexistence. Alternatively, using a symmetric version of the fitness function (3) with fluctuating fecundity, Schreiber et al. [2019] demonstrated numerically that the invasion growth rates *r*_*j*_ increase with positive autocorrelations. However, they did not analyze this numerical trend.

Autocorrelated fluctuations also can lead to an alternative stabilizing mechanism when (i’) there is a negative covariance between environment and competition, (ii) there are species-specific environmental responses, and (iii’) species are less sensitive to competition in years of good environmental conditions. Condition (iii’) arises when adult survival fluctuates. Condition (i’) occurs when fluctuations in survival are negatively auto-correlated as populations densities are higher in years following higher survival, but survival in the following year tend to be lower. Our analytical and numerical results show, however, that the strength of this stabilizing effect is weaker for a given magnitude of negative autocorrelation, than the strength of the storage effect for the same magnitude of positive autocorrelation. Negatively autocorrelated environments can arise in a variety of ways [Metcalf and Koons, 2007]. For example, approximately 5% of sites analyzed by [Sun et al., 2018] exhibit negatively autocorrelated rainfall. Interestingly, Adler and Levine [2007] found that species richness in central North American grasslands increased most in wet years that followed dry years. Alternatively, models and empirical studies show that overcompensatory or delayed density-dependent feedbacks can generate negatively autocorrelated fluctuations in densities [May, 1976, Tilman and Wedin, 1991, Crone and Taylor, 1996, Gilg et al., 2003]. If these fluctuations in densities occur in an herbivore, pathogen, or predator, then they can generate negatively autocorrelated fluctuations in survival of competing plants, hosts, or prey. Finally, competing species that exhibit two to three generations per year may experience negative autocorrelations in seasonal environments [Metcalf and Koons, 2007].

Autocorrelated fluctuations in survival or fecundity can generate alternative stable states and drive complex shifts in ecological outcomes. Alternative stable states arise as a stochastic priority effect: either species has a non-zero probability of being excluded, but the species at initially lower frequencies are more likely to be excluded. For small differences in mean fitness, stochastic induced alternative states arise whenever buffered populations experience a negative environment-competition covariance or whenever unbuffered populations experience a positive environment-competition covariance. Chesson [1988] highlights this possibility for annual plants with fluctuating seed survival, but his observation appears to be under-appreciated in the priority effects literature [Fukami, 2015, Fukami et al., 2016]. Shifts in temporal autocorrelations also generate complex shifts in ecological outcomes when one species has an inherent competitive advantage over the other, such as a higher mean environmental response or less variability in their environmental response. Under these circumstances, shifts from negative to positive temporal autocorrelations can result in shifts from a stochastic priority effect to competitive exclusion to coexistence (Fig. 3).

In conclusion, temporally autocorrelated environmental fluctuations indirectly generate a covariance between environmental conditions and the strength of competition. When this covariance is positive and there is population buffering this leads to a storage effect [Chesson, 1988, 1994]. As positive autocorrelations are seen in many climatic variables, accounting for these autocorrelations in data-driven models [Chu and Adler, 2015, Ellner et al., 2016, 2018] likely will lead to more empirically based examples of the storage effect. In contrast, when there is a negative environment-competition covariance, an alternative stabilizing mechanism to the storage effect arises provided species are less sensitive to competition in years where environmental conditions are favorable. As there are simple, ecologically plausible conditions that generate this alternative stabilizing mechanism, it will be exciting to see whether or not empirically based demonstrations will be found.

## Acknowledgments

Many thanks to Gyuri Barabasi, an anonymous reviewer, Kevin Gross, and Jennifer Lau for providing encouraging comments and exceptionally useful feedback on an earlier draft of the manuscript. The anonymous reviewer provided the excellent suggestion of applying Chesson’s community average mechanism to the covariance term (8). This research was supported in part by the U.S. National Science Foundation grant DMS-1716803.

## Appendix

In this appendix, I provide the mathematical details of the analysis of the deterministic and stochastic versions of the model (1) from the main text. Define the new coordinate system *x*_*i*_ = *a*_*i*_*n*_*i*_ in which (1) from the main text becomes

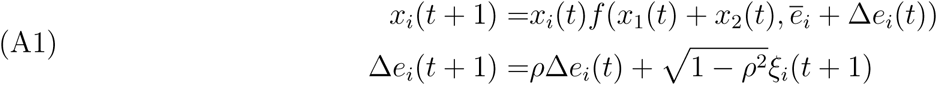

where *ξ*_*i*_(1), *ξ*_*i*_(2), … are a sequence of i.i.d. random variables with mean 0, variance 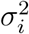 and cross-correlation *τ*. Throughout this analysis, I assume that *f* is a continuous, positive function, *C* ↦ *f* (*C, e*) is a decreasing function for all *e, e* ↦ *f* (*C, e*) is an increasing function for all *C*, and *C* ↦ *Cf* (*C, e*) is an increasing, bounded function. The first two assumptions ensure that fitness depends continuously on competition and environment, decreases with competition, and increases with the environmental variable. The third assumption corresponds to compensating density dependence and populations remaining bounded. A classic example of such a fitness function is the Beverton-Holt function with survival i.e *f* (*C, e*) = λ*/*(1 + *C*)+ *s* where either the maximal fitness λ of survivorship *s* are functions of *e*. Finally, I assume that *ξ*_*i*_(*t*) take values in a compact set and |*ρ*| < 1. Collectively, these assumptions imply that the dynamics of (A1) are dissipative i.e. there is a compact set in *K* ⊂ [0, ∞)^2^ ℝ^2^ such that non-negative solutions of (A1) eventually enter and remain in *K* for sufficiently large *t*.

Below, I first analyze the deterministic dynamics of (A1) i.e. when Δ*e*_*i*_(*t*) = 0 for all *t*. This analysis shows that there are three possible competitive outcomes: species 1 excludes species 2, species 2 excludes species 1, or neutral coexistence in which the dynamics converge to a line of equilibria. Next, I analyze the stochastic dynamics of (A1) in three steps. First, using results from Benaïm and Schreiber [2009], I provide a condition that ensures each species can persist in isolation. When this condition holds, each species’ dynamics converges to a unique stationary distribution 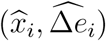. When this condition doesn’t hold, the species asymptotically converges to extinction with probability one. For the remainder of the stochastic analysis, I assume that the condition for each species persisting holds. Under this assumption, I use results from Benaïm and Schreiber [2019] to characterize coexistence and exclusion using the invasion growth rates *r*_*j*_. Finally, I use a diffusion type of approximation to derive the approximations for the invasion growth rates *r*_*j*_ presented in the main text.

### Deterministic Analysis

The model assumptions imply that (A1) is a strictly monotone, planar map with respect to the competitive ordering [see, e.g., Smith, 1998]. Each species persists individually if 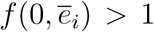 for *i* = 1, 2. Assume this condition holds for each species. Then, there are three cases to consider. First, assume that 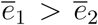. Define 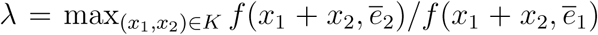 which is strictly less than 1 as 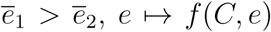 is a strictly increasing, continuous function, and *K* is compact. Given any solution (*x*_1_(*t*), *x*_2_(*t*)) to (A1) with *x*_1_(0) > 0, we get *x*_2_(*t*)*/x*_1_(*t*) ≤ λ^*t*^(*x*_2_(0)*/x*_1_(0)). As *x*_1_(*t*) is uniformly bounded i.e. (*x*_1_(*t*), *x*_2_(*t*)) ∈ *K, x*_2_(*t*) converges to 0 as *t* → ∞. Similarly, if 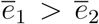, species 2 excludes species 1. Finally, consider the case that 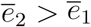. Then *x*_1_(*t*)*/x*_2_(*t*) = *x*_1_(0)*/x*_2_(0) for all time whenever *x*_2_(0) > 0. Namely, the relative frequency of either species doesn’t change over time. Moreover, as *y*(*t*) = *x*_1_(*t*) + *x*_2_(*t*) satisfies 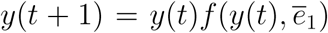 which a strictly monotone map with a unique positive equilibrium *y*^∗^ > 0, *x*_1_(*t*) + *x*_2_(*t*) converges to the *y*^∗^ as *t* → ∞. Hence, *x*_*i*_(*t*) converges to *x*_*i*_(0)*y*^∗^*/*(*x*_1_(0) + *x*_2_(0)) as *t* → ∞ for *i* = 1, 2. Thus, there is a globally stable line of equilibria given by {(*y*^∗^*p, y*^∗^(1 − *p*)) : 0 ≤ *p* ≤ 1}.

### General Stochastic Analysis

Define *g*(*x, e*) = log *f* (*x, e*) and assume |*ρ*| < 1. As the dynamics of Δ*e*_*i*_ are given by a multivariate autoregressive process where the linear term is contracting (i.e. |*ρ*| < 1) and *ξ*(*t*) are uniformly bounded, Δ*e*_*i*_(*t*) converge to a unique stationary distribution 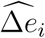 [see, e.g., Schreiber and Moore, 2018]. For each species *i* in isolation of the other species, their per-capita growth rate at low densities is 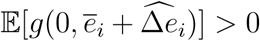. When this invasion growth rate is positive and *x*_*i*_(0) > 0, Theorem 1 in [Benaïm and Schreiber, 2009] implies that solutions (*x*_*i*_(*t*), Δ*e*_1_(*t*), Δ*e*_2_(*t*)) for the species *i* subsystem with *x*_*i*_(0) > 0 converges to a unique stationary distribution 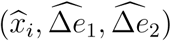. In contrast, if 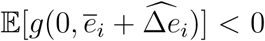, Proposition 1 in [Benaïm and Schreiber, 2009] implies that *x*_*i*_(*t*) converges to 0 with probability one. From the rest of the stochastic analysis, assume that 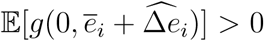 for both species *i* = 1, 2.

The invasion growth rate of species *j* ≠ *i* when species *i* is at its stationary distribution equals

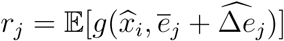

Theorem 1 from [Benaïm and Schreiber, 2019] implies that the two species coexist (in the sense of stochastic persistence) whenever *r*_1_ > 0 and *r*_2_ > 0. In contrast, if *r*_*j*_ < 0, then Theorem 3 from [Benaïm and Schreiber, 2019] implies that species *j* goes extinct with high probability whenever its initial density is sufficiently low. Under suitable accessibility assumptions (see Benaïm and Schreiber [2019] for definitions), stronger conclusions hold when *r*_*j*_ < 0. In particular, if *r*_1_ > 0 > *r*_2_ (respectively, *r*_2_ > 0 > *r*_1_), then species 1 excludes species 2 with probability one (respectively, species 2 excludes species 1) whenever *x*_1_(0) > 0 (respectively, *x*_2_(0) > 0). Alternatively, if *r*_1_ < 0 and *r*_2_ < 0, then either species, with complementary positive probabilities, goes extinct whenever both are initially present i.e. a stochastic priority effect.

### Diffusion scaling to approximate *r*_*j*_

For any *ε* > 0, assume that (i) *ξ*_*i*_(*t*) = *εη*_*i*_(*t*) where *η*_*i*_(*t*) are i.i.d. with mean zero, variance *v*_*i*_, and 𝔼[*η*_1_(*t*)*η*_2_(*t*)] = *τ*, and (ii) 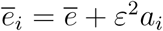. To ensure each species persists in the absence of the other, assume that the invasion growth rate 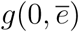 is positive. Then there exists *x*^∗^ > 0 be such that 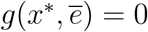 corresponds to the single species, equilibrium density in the absence of fitness differences and environmental fluctuations. Assume that *g*(*x, e*) is three times continuously differentiable and let *g*_*x*_, *g*_*e*_, *g*_*xe*_, etc. denote the partial derivatives 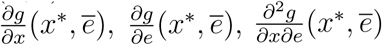, etc. evaluated at 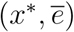.

Consider species *i* as the common or resident species and *j* ≠ *i* as the rare or invading species. Proposition 1(iii) from [Benaïm and Schreiber, 2019] implies that the invasion growth of 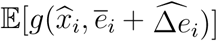, at its stationary distribution equals zero. Thus, taking a Taylor’s expansion and defining 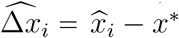 yields

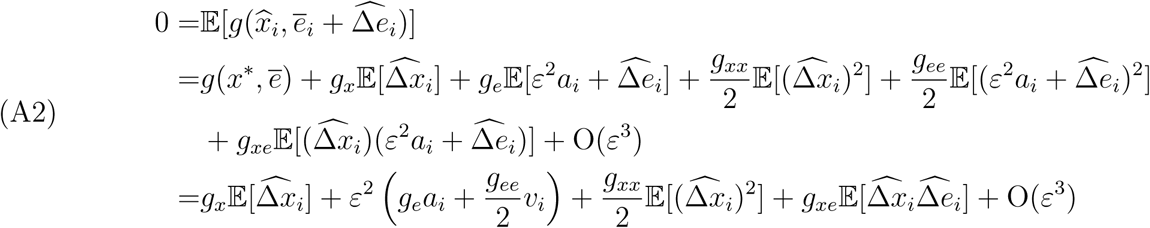

as 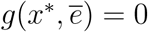 due to the definition of 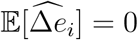, and 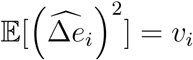, and 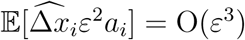. Similarly, we get

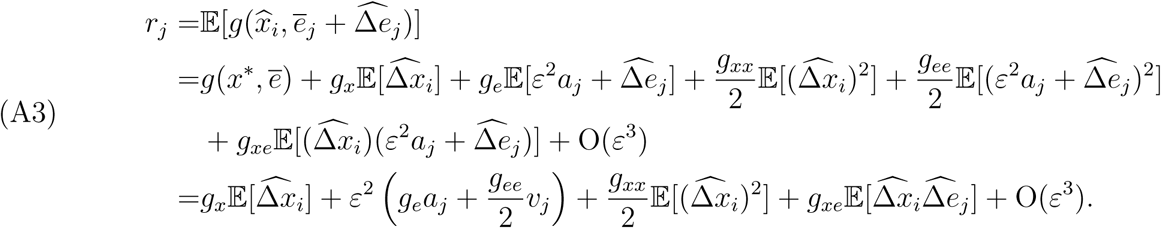

Subtracting equation (A2) from equation (A3) yields

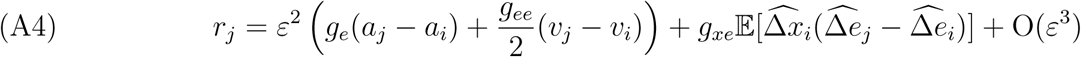

To get an explicit expression for 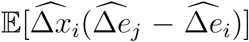, one can approximate the dynamics of *x*_*i*_ as a first order autoregressive process by linearizing (A1) at 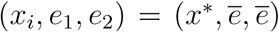. To this end, define *F* (*x, e*) = *xf* (*x, e*) and *F*_*x*_, *F*_*e*_ as the partial derivatives 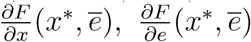. Then, the first-order autoregressive approximation of (A1) is

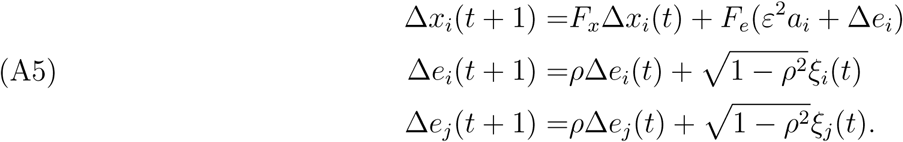

Equivalently, defining *z* = (Δ*x*_*i*_, Δ*e*_*i*_, Δ*e*_*j*_)

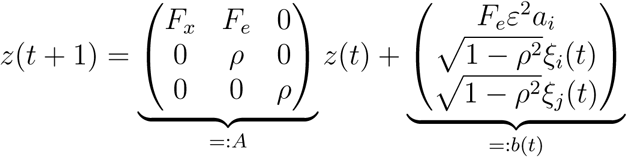

The covariance matrix of *Z*(*t*),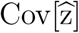, at stationarity [see, e.g., Schreiber and Moore, 2018] satisfies

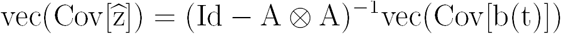

where vec(∗) is a column vector given by concatenating the columns of its argument ∗ and ⊗ denotes the Kroenker product. Carrying out this calculation yields the approximations

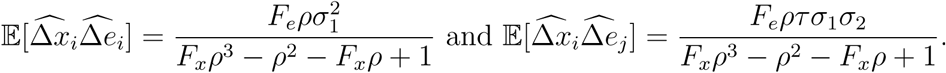

To see that the denominator of these expressions is positive for |*ρ*| < 1, define *h*(*y*) = *yρ*(*ρ*^2^ 1) *ρ*^2^ +1. As *F*_*x*_ ∈ (0, 1) (i.e. the equilibrium is stable as the dynamics are compensatory), one needs to only consider *h*(*y*) for *y* ∈ (0, 1). The minimum of *h*(*y*) occurs either at *y* = 1 or *y* = −1 or at 0 < *y* < 1 where *h*′(*y*) = 0. For |*ρ*| < 1, *h*′(*y*) = *ρ*(*ρ*^2^ − 1) = 0 only when *ρ* = 0 in which case *h*(*y*) = 1 > 0. As *h*(0) = 1 − *ρ*^2^ > 0 for |*ρ*| < 1 and *h*(1) = *ρ*^3^ − *ρ*^2^ − *ρ* + 1 > 0 for |*ρ*| < 1, it follows that *h*(*y*) > 0 for *y* ∈ (0, 1) whenever |*ρ*| < 1.

Define

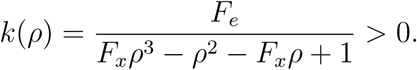

Then,

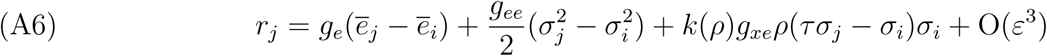

as claimed in the main text. Let *H*(*ρ*) = *ρk*(*ρ*). Then

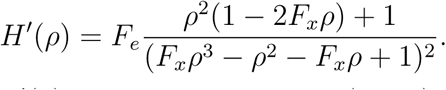

As 0 < *F*_*x*_ < 1, the numerator of *H*′(*ρ*) is positive for all *ρ* ∈ (−1, 1). Therefore, as claimed in the main text, *H*(*ρ*) is an increasing function of *ρ*. Finally, consider *ρ* > 0. Then *k*(−*ρ*) < *k*(*ρ*) if and only if *F*_*x*_*ρ*^3^ − *ρ*^2^ − *F*_*x*_*ρ* + 1 < −*F*_*x*_*ρ*^3^ − *ρ*^2^ + *F*_*x*_*ρ* + 1. This occurs if and only if *ρ*^3^ < *ρ* which is true whenever *ρ* ∈ (0, 1). Thus, |*ρk*(*ρ*)| > | − *ρk*(−*ρ*)| for any *ρ* ∈ (0, 1) as claimed in the main text.

